# TGF-β2/OPTN/FOXC1/*miR-200* axis regulates intraocular pressure dynamics in trabecular meshwork cells

**DOI:** 10.1101/2024.06.06.593675

**Authors:** Chenna Kesavulu Sugali, Navya Naidu Gajula, Suresh Chava, Aramati Bindu Madhava Reddy

## Abstract

Glaucoma is the second leading cause of irreversible blindness globally, with elevated intraocular pressure (IOP) being its primary risk factor. Current therapeutic approaches, such as beta-blockers, alpha-adrenergic agonists, Rho-kinase inhibitors, etc., aim to reduce IOP levels. However, the molecular mechanisms underlying altered IOP remain poorly understood. In this study, we have treated primary human trabecular meshwork cells (HTM) with exogenous dexamethasone (dex) or transforming growth factor beta-2 (TGF-β2) to investigate its effects on glaucoma candidate genes. Interestingly, our findings reveal that FOXC1 acts as a repressor to *CYP1B1*, and optineurin (OPTN) facilitates the ubiquitination of FOXC1, thereby inducing CYP1B1 expression. Further, we discovered that the *miR-200* family and other miRNAs regulate these glaucoma-candidate genes. Furthermore, TGF-β2 downregulates the *miR-200* family, whereas the *miR-200* family targets *FOXC1*, exerting reversible effects by altering the extracellular matrix. FOXC1 positively regulates *CLOCK*, one of its target genes. Besides, CLOCK/BMAL1 has binding sites on miR-200 family promoters. Modulating the TGF-β2/OPTN/FOXC1/*miR-200* axis appears critical in regulating IOP dynamics through CLOCK/BMAL1-mediated daily rhythmicity in the anterior segment of the eye.

**Graphical Abstract:** 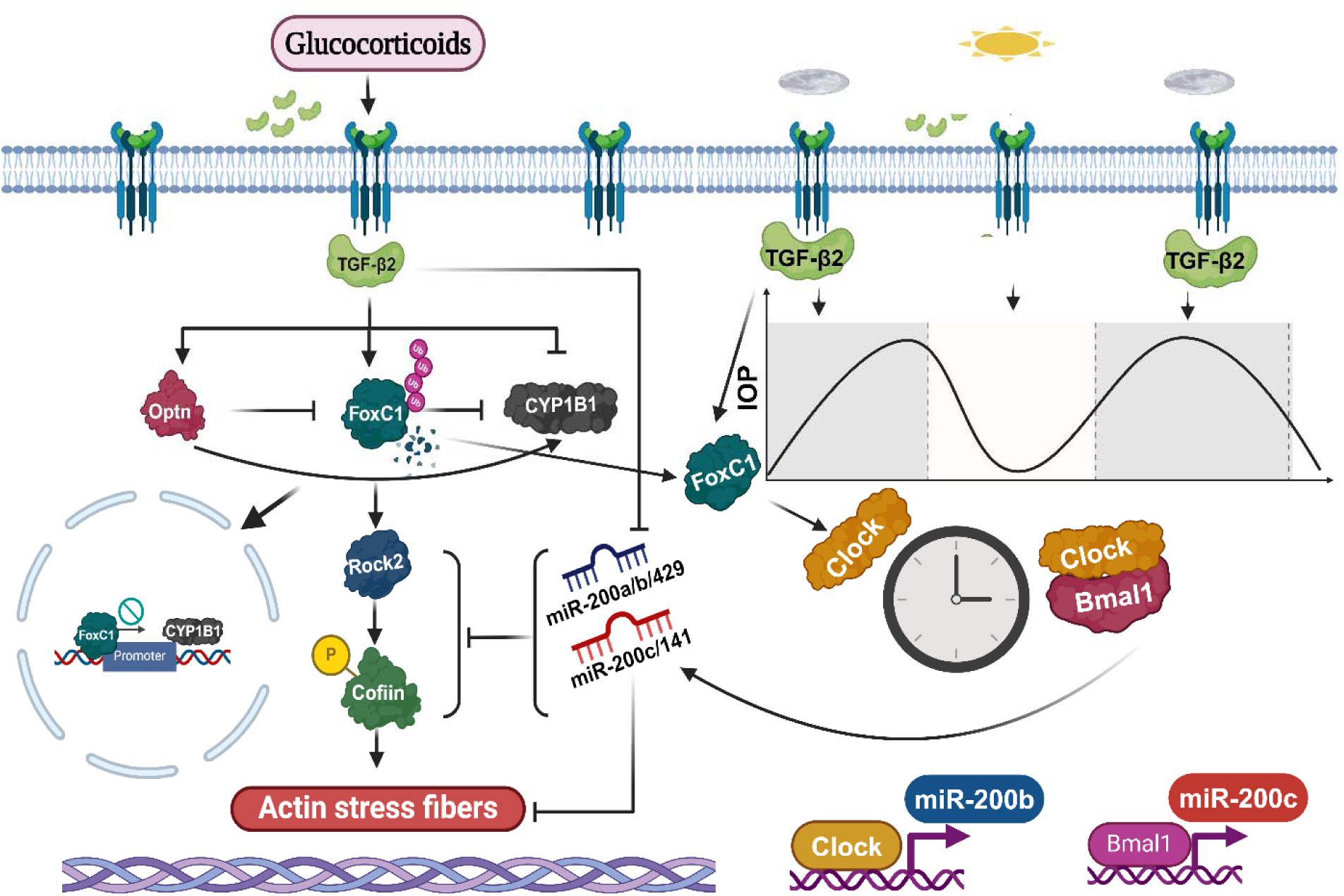

## 1. Introduction

Glaucoma is the leading cause of blindness after cataracts ^1–3^. The primary risk factor is elevated IOP ^4^ due to hindered drainage of aqueous humor (AH) through the trabecular meshwork (TM), a tissue located in the iridocorneal angle in the anterior chamber of the eye ^5^. Aberrant synthesis and deposition of ECM proteins like fibronectin, collagen, elastin, laminins, glycosaminoglycans, and periostin in the TM lead to increased outflow resistance ^6–10^. Studies have shown that TM contractility hampers aqueous humor (AH) drainage whereas, TM relaxation facilitates it ^5,11^. Consequently, preserving the functional integrity of the cytoskeleton within TM tissue is critical for the AH outflow pathway and the regulation of ocular hypertension ^12^.

Glaucoma is a complex disease associated with mutations in various genes, including those encoding cytochrome P4501B1 (CYP1B1) enzyme ^13,14^, myocilin (MYOC) ^15^, optineurin (OPTN) ^16^, forkhead box C1 (FOXC1) ^15,17^, latent transforming growth factor beta binding protein 2 (LTBP2) ^18^, and others, linked to its onset or progression ^19^. However, little is known about these proteins’ role in regulating the eye’s outflow pathways. Studies have indicated that elevated levels of cytokines, such as TGF-β2 in the aqueous humor of glaucoma patients, can induce the expression of extracellular matrix (ECM) proteins like fibronectin, collagen, laminin, and glycosaminoglycan, as well as cytoskeletal reorganization by promoting the formation of actin stress fibers within the TM, which could lead to TM fibrosis ^20–27^. These alterations result in increased rigidity and stiffness of TM cells, hindering aqueous outflow and increasing IOP ^21,28^. However, the precise mechanisms underlying the regulation of TGF-β2 and glaucoma candidate genes (*CYP1B1, MYOC, OPTN, FOXC1,* and *LTBP2*) during IOP buildup remain unclear.

Since different classes of genes come together in mediating the disease, regulatory factors like miRNAs, that target multiple genes play a crucial role in the pathogenesis of glaucoma. Microarray and transcriptome studies have uncovered distinct miRNA expression patterns associated with glaucomatous changes in TM integrity, AH outflow regulation, and the fate of retinal ganglion cells (RGCs) ^29–37^. Recent reports have specifically highlighted the differential expression of *miRNA-29b* ^38,39^, *miR-18a* ^41^, *miRNA-204* ^40^, *miRNA-200c* ^42^ and miRNA cluster, *miR-143/145* ^43^ in glaucomatous TM and AH, suggesting that miRNAs may modulate TM contraction and IOP. However, conclusive genetic evidence demonstrating miRNA involvement in IOP regulation is still lacking.

Our findings demonstrate that the glucocorticoids dexamethasone (dex) and prednisolone (pred) as well as the cytokine, TGF-β2 regulate the expression of glaucoma candidate genes (*CYP1B1, MYOC, OPTN, FOXC1,* and *LTBP2*). We observed a dynamic interplay among CYP1B1, FOXC1, and OPTN within the TM, affecting IOP regulation. Dysregulation of these candidate genes alters the Rho-Rock pathway and actin stress fiber formation. Furthermore, TGF-β2 downregulates the expression of putative miRNAs. We also demonstrate that the *miR-200* family can inhibit TGF-β2 and ROCK2 expression in TM cells, thus preventing actin stress fiber formation and promoting TM relaxation. This study unveils, for the first time, the significance of the TGF-β2/OPTN/FOXC1/*miR-200* axis in ocular hypertension, TGF-β2 secretion, and TM contraction, offering novel insights into glaucoma pathogenesis.

## 2. Results

### 2.1 Glucocorticoids and TGF-β2 can regulate the expression of glaucoma candidate genes in primary HTM cells

Previous studies have established a correlation between elevated IOP and the use of topical corticosteroids, such as dex and pred ^44,45^. For instance, dex can induce *TGF-*β*2* expression in HTM cells ^46,47^. Moreover, genes like *TGF-*β*2*, *CYP1B1*, *FOXC1*, *OPTN*, *MYOC,* and *LTBP2* have been associated with glaucoma development ^48,49^. Based on these findings, we sought to investigate the regulatory effects of glucocorticoids on the expression of these glaucoma-associated genes. Our results demonstrate that the addition of dex and pred led to an upregulation of MYOC in HTM cells, consistent with previous findings (Figures 1A, B) ^50^. Furthermore, glucocorticoid treatment resulted in the downregulation of the protein and mRNA levels of CYP1B1 (Figures 1A, C), and the upregulation of FOXC1 and OPTN at both the protein and mRNA levels (Figures 1A, D-E). Similarly, glucocorticoid treatment increased LTBP2 expression (Figure 1F). Additionally, we observed an increase in the levels of active TGF-β2 and its expression in response to dex and pred, as depicted in (Figures 1G, H). Further, treatment with dex for 3 weeks elevated IOP over time in the eyes of C57BL/6J mice (Figure 1I). Interestingly, these mice exhibited noticeable changes in optic disc cupping following treatment (Figure 1J). Moreover, this treatment enhanced the accumulation of TGF-β2 in the anterior chamber of the eye of C57BL/6J mice, as evidenced by immunohistochemistry (Figure 1K). These findings suggest that glucocorticoids can alter the expression of glaucoma candidate genes apart from inducing TGF-β2 production. However, it isn’t clear if the genes are regulated through TGF-β2. To elucidate the molecular mechanisms by which TGF-β2 mediates its effect, primary HTM cells were treated with TGF-β2 for 48h, and cell lysates were analyzed for protein and RNA expression. Similar to dex, we observed upregulation of MYOC at both the protein and mRNA levels (Figures 1L, M), CYP1B1 was downregulated both at both the protein and mRNA levels in response to TGF-β2 in HTM cells (Figures 1L, N) while FOXC1, OPTN, LTBP2 were upregulated (Figures 1L, O-Q). In summary, our results indicate that both glucocorticoids and TGF-β2 can influence the expression of glaucoma candidate genes in trabecular meshwork cells.

**Figure 1:**
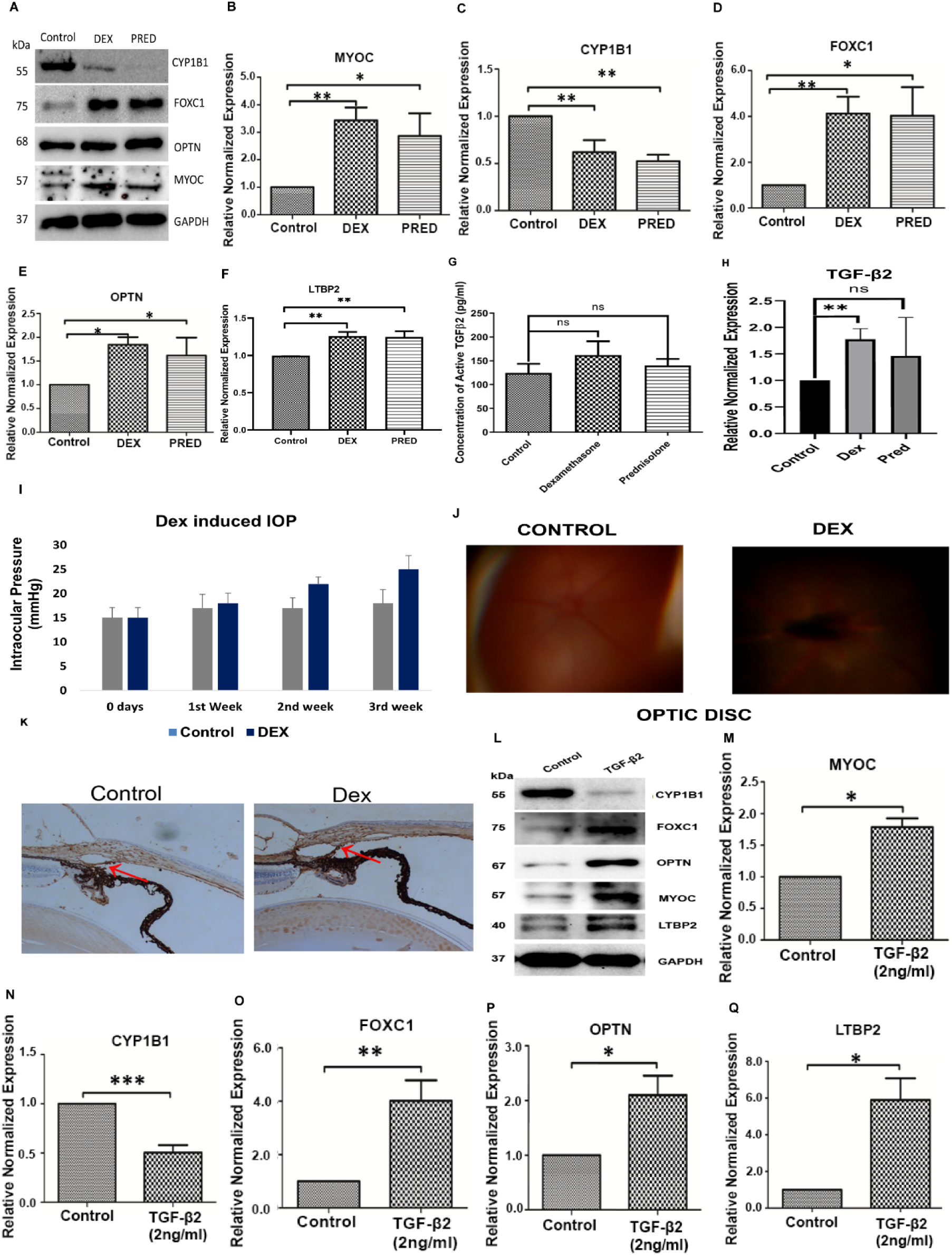
Glucocorticoids and TGF-β2 regulate the expression of glaucoma candidate genes in primary HTM cells. (A) Western blot analysis in HTM cells 48h post-treatment of dex and pred. (B-F) RT-PCR analysis 48h post-treatment of dex and pred. (G) ELISA 48h post-treatment of dex and pred. (H) RT-PCR analysis of TGF-β2 48h post-treatment of dex and pred. (I) IOP measured every week up to 3 weeks post-treatment of dex in C57BL/6J mice. (J) Optic disc images captured after 3 weeks post-treatment of dex in C57BL/6J mice. (K) IHC of TGF-β2 3 weeks post-treatment of dex in C57BL/6J mice. (L) Western blot analysis in HTM cells 48h post-treatment of TGF-β2. (M-Q) RT-PCR analysis 48h post-treatment of TGF-β2. Bars represent the standard deviation in three different experiments. Asterisk (*) (**) (***) represents significance at p<0.5 p<0.05 p<0.005 respectively between treatments and control. Standard deviation and significance were calculated by using GraphPad Prism 8.0.2.

### 2.2 CYP1B1, FOXC1, and OPTN interregulate each other in primary HTM cells

To investigate how FOXC1, CYP1B1, and OPTN regulate IOP and contribute to glaucoma development, we employed siRNA knock-down and ectopic overexpression to manipulate the expression of *CYP1B1*, *FOXC1*, and *OPTN* genes. We then assessed the interplay among these genes using western blot and RT-PCR. Our data revealed that *FOXC1* knockdown led to an increase in CYP1B1 expression (Figures 2A, D). Conversely, *CYP1B1* knockdown did not significantly influence FOXC1 (Figures 2B, D) and OPTN (Figures 2C-D) expression. While *FOXC1* ectopic expression decreased CYP1B1 expression (Figure 2E). Manipulating *FOXC1* levels did not affect OPTN expression in primary HTM cells, on the other hand (Figure 2C, E). These results suggest that the transcription factor FOXC1 regulates CYP1B1 expression in TM cells (Figures 2A, 2D-E).

**Figure 2:**
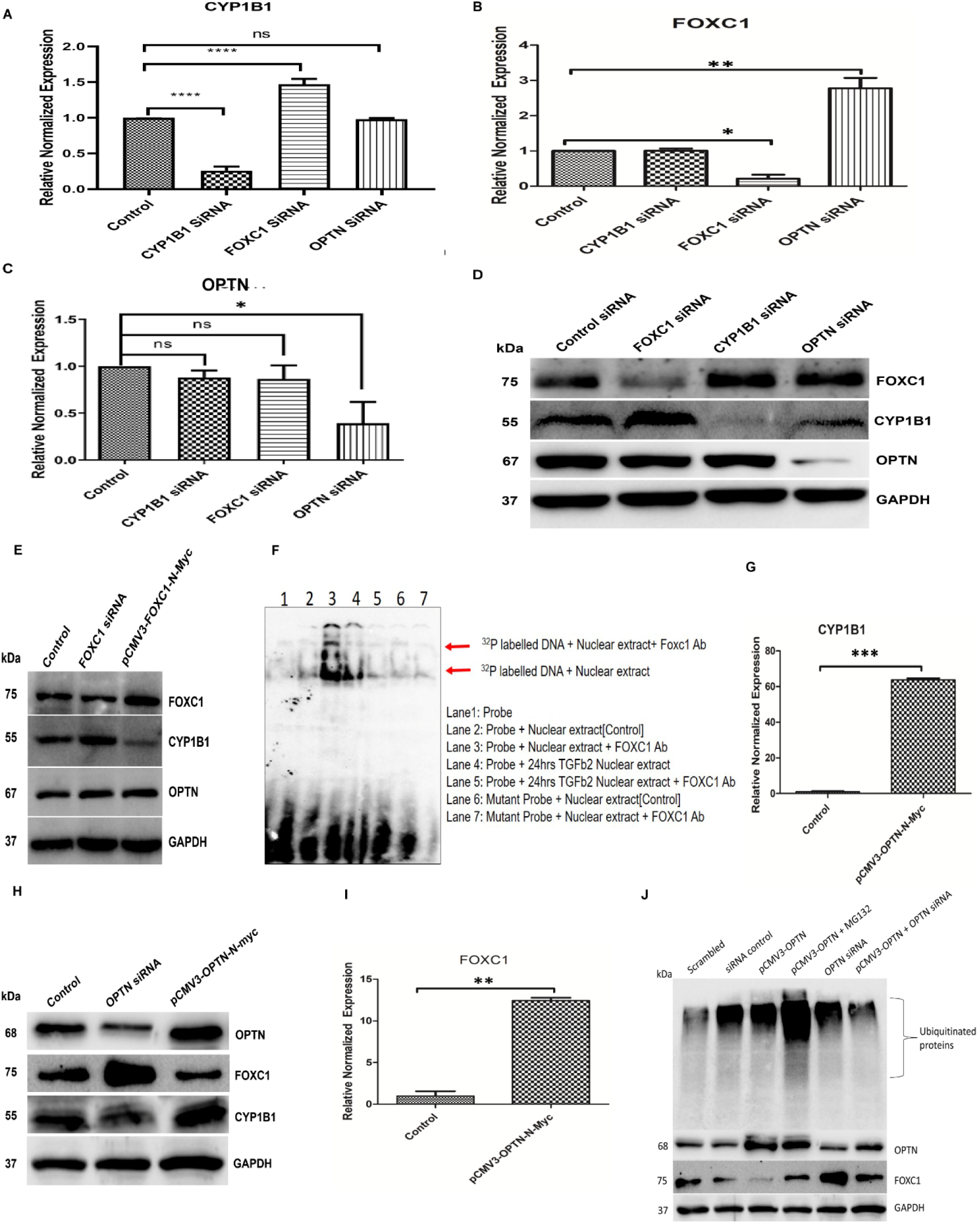
CYP1B1, FOXC1, and OPTN interregulate each other in primary HTM cells. (A-C) RT-PCR analysis 48h post-transfection of respective siRNAs. (D) Western blot analysis upon knock-down of *CYP1B1*, *FOXC1,* and *OPTN* for 48h. (E) Western blot analysis upon knock-down and over-expression of *FOXC1* for 48h. (F) EMSA showing the binding of FOXC1 onto *CYP1B1* promoter in HTM cells. (G, I) RT-PCR analysis upon overexpression of OPTN for 48h in HTM cells. (H) Western blot analysis upon overexpression and knockdown of OPTN in HTM cells. (J) Western blot analysis hinting to ubiquitination of FOXC1, by overexpression and knock-down of *OPTN* in combination with treatment of MG132.

We subsequently investigated whether FOXC1 can bind to the *CYP1B1* promoter. Sequence analysis revealed a FOXC1 consensus binding site (TAAACAACC) at positions –2622 to –2631 of the CYP1B1 (GenBank-U56438.1) promoter. To characterize and confirm the transcriptional binding efficiency of FOXC1 to the *CYP1B1* promoter, we performed an electrophoretic mobility shift assay (EMSA) using 21-bp oligos of the region containing the *CYP1B1* consensus sequence with and without sequence variations. EMSA analysis using nuclear fractions extracted from primary HTM cells revealed that FOXC1 binds to the *CYP1B1* promoter, as confirmed by mutant probes with and without super shift with FOXC1 antibody (Figure 2F). This finding agrees with our observation that ectopic *FOXC1* overexpression decreases CYP1B1 (Figure 2E), thus confirming that TGF-β2-induced FOXC1 represses CYP1B1 in HTM cells (Figure 2F).

Our results also demonstrated that *OPTN* knockdown did not impact CYP1B1, but significantly increased FOXC1 expression (Figure 2A, B). Moreover, ectopic overexpression of *pCMV3-OPTN* significantly upregulated CYP1B1 (Figure 2G-H) mRNA levels and downregulated FOXC1 expression at both the protein and mRNA levels (Figure 2I), while reducing FOXC1 protein levels (Figure 2J). This observation suggests that FOXC1 may undergo ubiquitination in the presence of OPTN in primary HTM cells. To verify this, we treated TM cells expressing ectopically overexpressed *OPTN* with and without MG132 (an inhibitor of the proteasomal pathway) or *OPTN* siRNA. We observed that *OPTN* siRNA or MG132 prevented FOXC1 ubiquitination despite *OPTN* overexpression (Figure 2J). Taken together, our results suggest that CYP1B1, FOXC1, and OPTN form a coordinated regulatory axis in primary HTM cells.

### 2.3 Loss of OPTN induces TGF-β2 and activates the Rho/ROCK signaling pathway in primary HTM cells

Our data indicates that TGF-β2 can induce OPTN, which subsequently regulates the expression of CYP1B1 and FOXC1. However, the interplay between these factors in the pathophysiology of glaucoma remains poorly understood. Therefore, we sought to investigate the role of OPTN in regulating TGF-β2 in primary HTM cells. We transfected primary HTM cells with either *pCMV3-OPTN* or *OPTN* siRNA, along with appropriate controls (pCMV3 empty vector or Allstar siRNA negative control) to modulate *OPTN* expression ectopically. Following experimental treatments, we collected and analyzed conditional medium, RNA, and protein for TGFβ2 levels using qRT-PCR, ELISA, or western blot, respectively.

Our findings revealed that *OPTN* knockdown increased TGF-β2 levels, whereas overexpression of *OPTN* was associated with a reduction in TGF-β2 levels (Figure 3(I) A, B). Given that *OPTN* loss induces TGF-β2 expression, we further examined the Rho/ROCK signaling, a downstream effector pathway of TGF-β2 implicated in ECM remodeling and the outflow pathway in TM tissue. Interestingly, *OPTN* knockdown resulted in the upregulation of both ROCK2 and p-cofilin (Figure 3(I) C), confirming that *OPTN* loss can induce TGF-β2 expression and activate the downstream Rho/ROCK signaling pathway.

**Figure 3(I):**
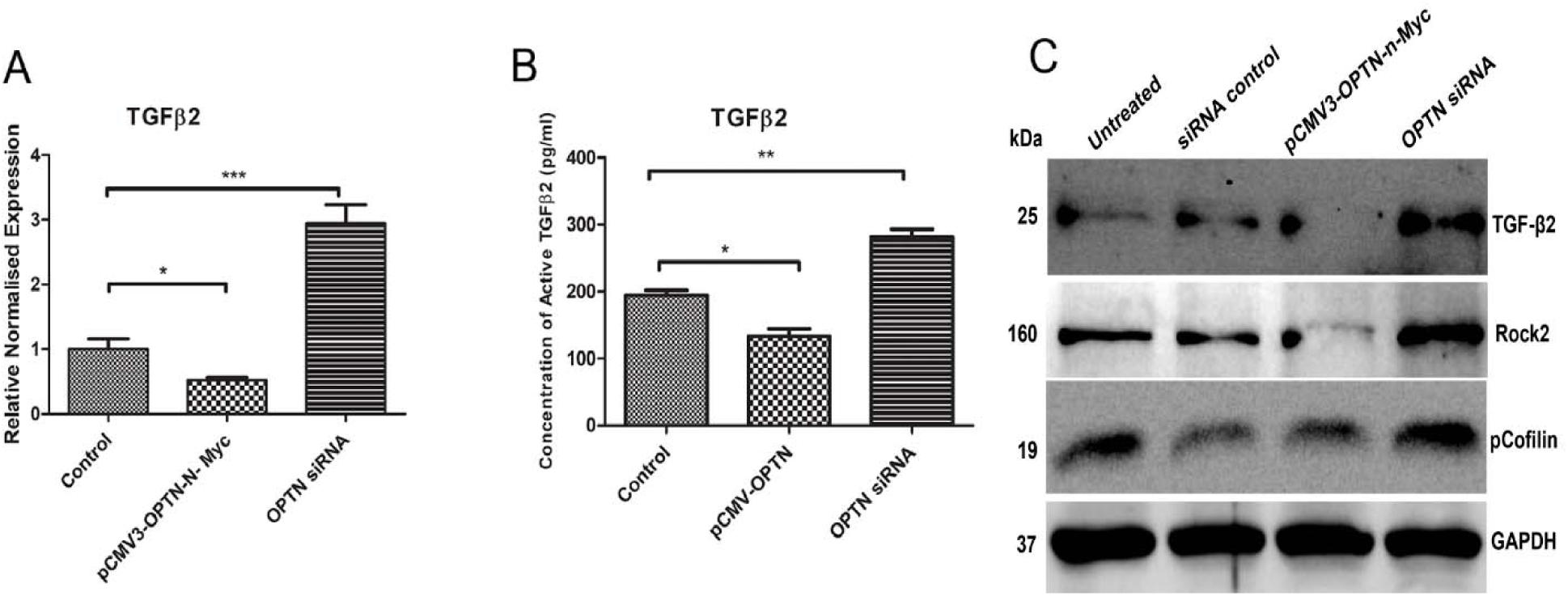
Loss of OPTN induces TGF-β2 and activates the Rho/ROCK signaling pathway in primary HTM cells. (A) RT-PCR analysis post 48h overexpression and knockdown of *OPTN*. (B) ELISA upon overexpression and knockdown of *OPTN* for 48h. (C) Western blot analysis of TGF-β2 induced ROCK pathway upon overexpression and knockdown of *OPTN* for 48h in HTM cells.

### 2.4 Knockdown of glaucoma candidate genes induces actin stress fibers and focal adhesions in the primary HTM cells

The regulation of Rho/ROCK signaling is critical in the formation of actin stress fibers, which contribute to increased resistance to outflow ^51,52^. To assess the role of *FOXC1*, *OPTN*, and *CYP1B1* in actin cytoskeletal reorganization, we used specific siRNAs to knock down their respective genes in primary HTM cells. We found that the loss of *FOXC1*, *OPTN*, and *CYP1B1* can induce the formation of actin stress fibers and focal adhesions in primary HTM cells compared to control cells. Additionally, TGF-β2-induced actin stress fibers served as a positive control (Figure 3(II)). Similar results were observed following treatment with 2,4,3’,5’-tetramethoxystilbene (TMS), a known chemical inhibitor of CYP1B1. Conversely, treatment with 2,3,7,8 Tetrachlorodibenzo-p-dioxin (TCDD), a known chemical inducer of CYP1B1, resulted in the suppression of stress fiber formation (Figure 3(II)).

**Figure 3(II):**
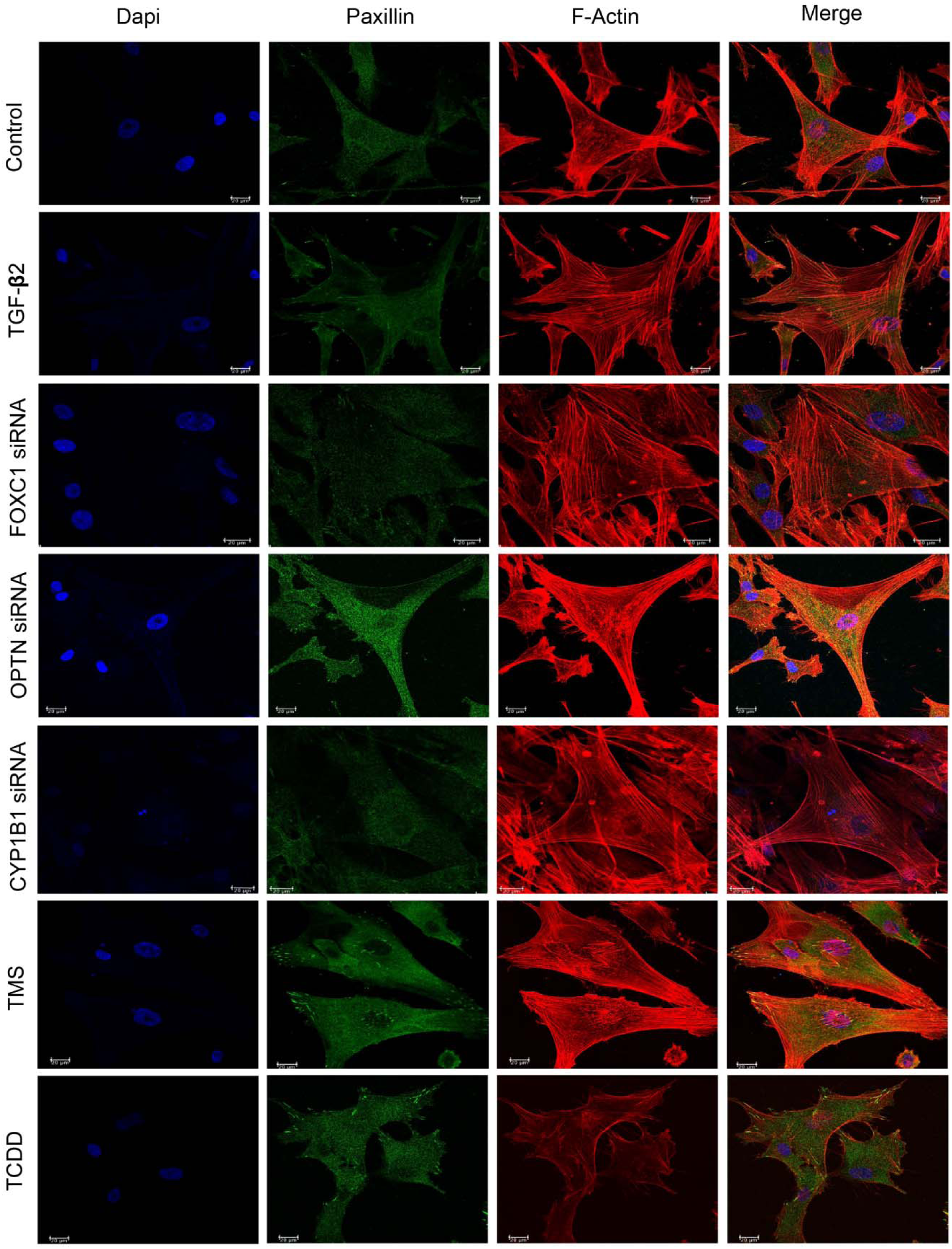
Knockdown of glaucoma candidate genes induces actin stress fibers and focal adhesions in the primary trabecular meshwork cells. Confocal imaging of HTM cells stained for focal adhesion protein, Paxillin (Green), and F-actin (Red) under respective conditions.

### 2.5 Ectopic expression of *miR-200* family, *miR-590*, *miR-496*, and *miR-548c* alters the expression of glaucoma candidate genes in primary HTM cells

Gene expression is tightly regulated by various mechanisms, including transcription and translation. miRNAs play a key role in modulating gene expression in eukaryotic cells ^53^. They are also involved in the pathogenesis of glaucoma and may influence the biological processes of genes associated with this condition ^31,32^. Our findings suggest that the loss of glaucoma candidate genes may impact the function of the trabecular meshwork. This prompted us to investigate whether these genes undergo post-transcriptional modifications by miRNAs. We identified several miRNAs, including *miR-200a*, *miR-200b*, *miR-200c*, *miR-429*, *miR-141*, *miR-590*, *miR-496*, and *miR-548c* (Table 1), which potentially target multiple glaucoma candidate genes (*CYP1B1*, *FOXC1*, *OPTN*, *MYOC*, and *LTBP2*), for further analysis using few bioinformatics tools. Our findings suggest that FOXC1 may negatively regulate CYP1B1. To investigate whether *FOXC1* is a direct target of the screened miRNAs, we transfected these miRNAs, along with *FOXC1* 3’UTR constructs cloned in the pMIR-Report vector, into HEK293T cell lines. After 24 hours, we conducted a dual luciferase assay and observed decreased luminescence in the *miR-200* family and *miR-548c* compared to the control (Figure 4A).

**Figure 4:**
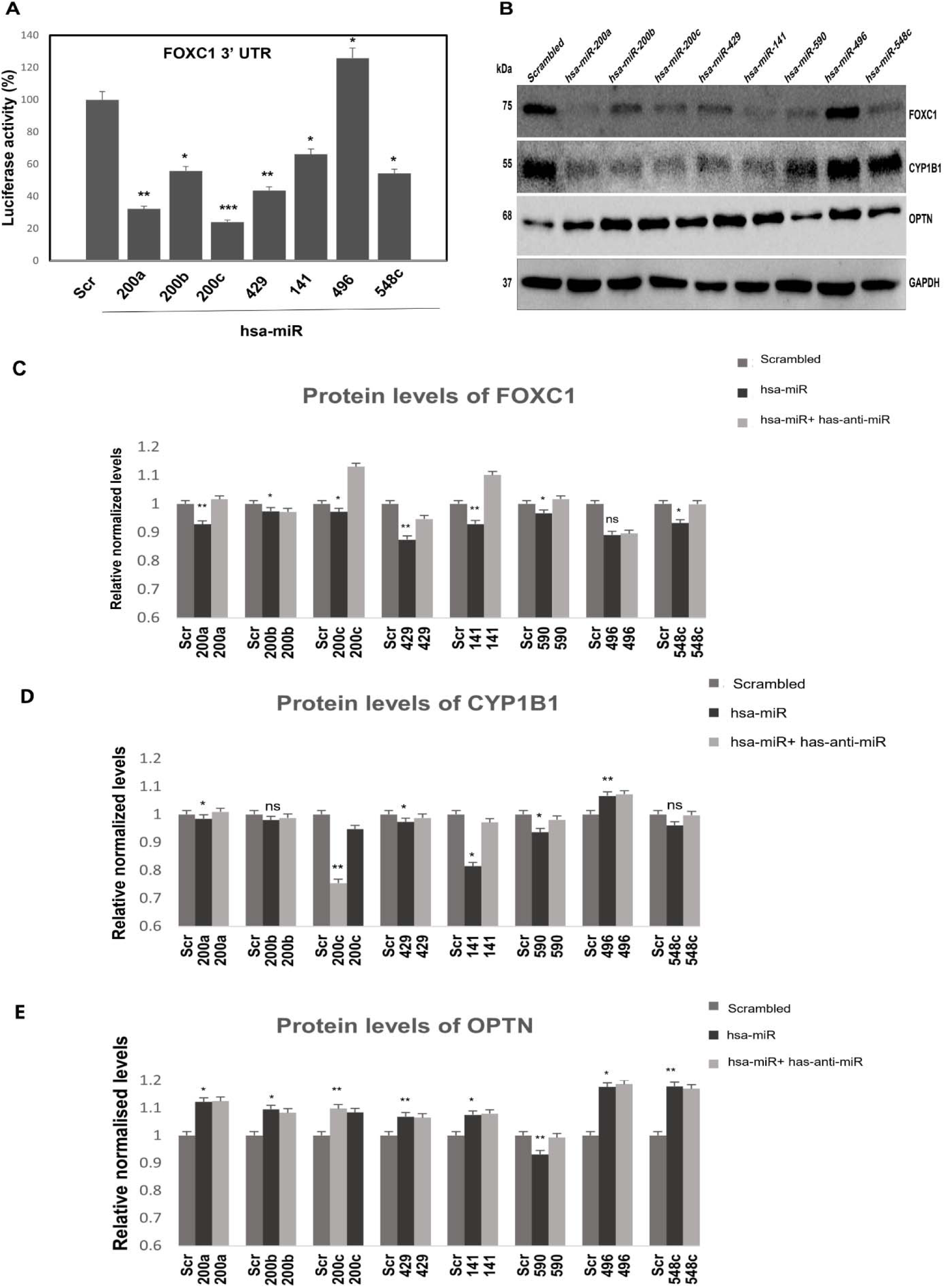
Ectopic expression of *miR-200* family, *miR-590*, *miR-496*, and *miR-548c* alters the expression of glaucoma candidate genes in primary HTM cells. (A) Dual luciferase assay in HEK293T cell lines harvested after 24h. Bars represent the standard deviation in three different experiments. (B) Western blot analysis 48h post-transfecting HTM cells with miRNAs. (C-E) Western blot quantification data of protein lysates upon transfection of respective miRs and anti-miRs.

In addition, we transfected primary HTM cells with miRNA mimics, with or without anti-miRs for 48h, followed by analysis of protein lysates. Western blot analysis revealed that the *miR-200* family, *miR-590,* and *miR-548c* downregulated the expression of FOXC1 and CYP1B1 (Figure 4B) whereas, anti-miRs targeting *miR-200a*, *miR-200c*, *miR-429*, and *miR-590* rescued their expression (Figure 4C, D). Furthermore, *miR-590* downregulated OPTN, while anti-miR targeting *miR-590* rescued its expression (Figure 4E). In summary, our results indicate that *FOXC1* seems to be a direct target for the *miR-200* cluster and *miR-548c*. Moreover, OPTN, along with CYP1B1 and FOXC1, appears to be targeted by certain miRNAs in HTM cells.

### 2.6 *miR-200* family, *miR-590,* and *miR-548c* inhibited TGF-β2 induced actin stress fibers and focal adhesion in primary HTM cells

TGF-β2 can induce the formation of actin stress fibers and focal adhesion in primary HTM cells ^21,28,54–56^, which correlates with increased outflow resistance in glaucoma. Our experiments suggest that certain miRNAs can target and downregulate the expression of TGF-β2 and ROCK2. To assess the formation of actin stress fibers and focal adhesion in primary HTM cells, we treated primary HTM cells with or without TGFβ2, along with and without miRNAs for 48h. Subsequently, we fixed the cells with Rhodamine phalloidin for actin stress fibers and paxillin for focal adhesions. We observed a decrease in actin stress fibers and paxillin (focal adhesion protein) in HTM cells following transfection with miRNAs, even in the presence of TGFβ2, compared to cells treated with TGFβ2 alone (Figure 5).

**Figure 5:**
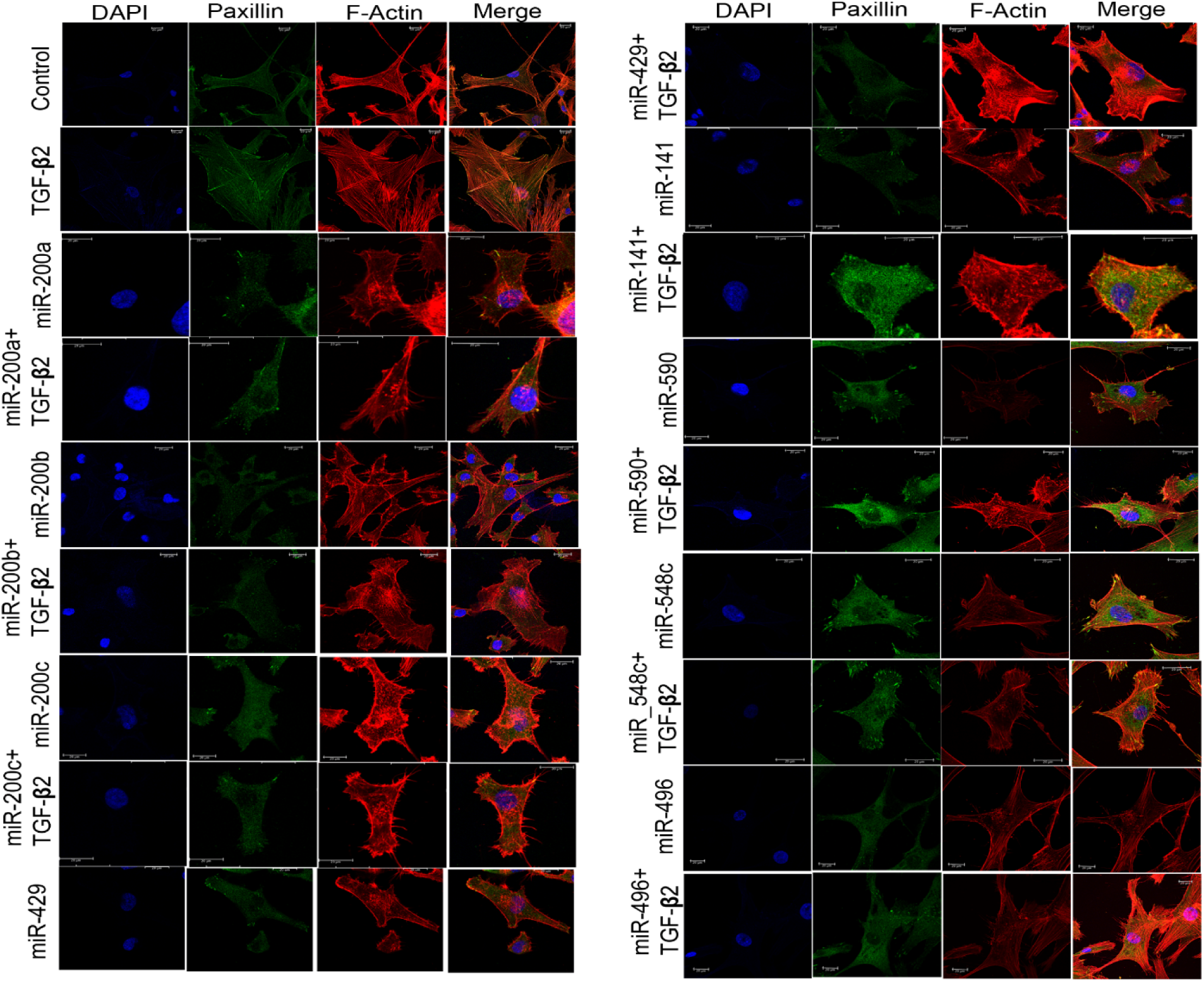
*miR-200* family, *miR-590* and *miR-548c* inhibited TGF-β2 induced actin stress fibers and focal adhesions in primary HTM cells. Confocal imaging of HTM cells stained for focal adhesion protein, Paxillin (Green) and F-actin (Red).

### 2.7 *miR-200* family, *miR-429*, *miR-590*, *miR-496* and *miR-548c* regulated contraction and relaxation of primary HTM cells

To evaluate the involvement of predicted miRNAs in the contraction and relaxation of HTM cells, we transfected primary HTM cells with and without these miRNAs for 48h. Subsequently, we embedded the cells in collagen gels for up to 24 hours. Following this, the cells were detached from the walls and treated with and without TGF-β2 (2ng/ml). We monitored TM cell contraction after 24h, and the extent of contraction was measured using the NIKON COOLPIX camera at the endpoint.

Our findings indicate that TGF-β2 induced the contraction of collagen-embedded primary HTM cells, whereas treatments with respective miRNAs significantly attenuated the TGF-β2-induced contraction of HTM cells embedded in collagen gels (Figure 6A). These results provide further evidence that the putative miRNAs can counteract TGF-β2-induced actin stress fiber formation, focal adhesion, and contraction of HTM cells.

**Figure 6:**
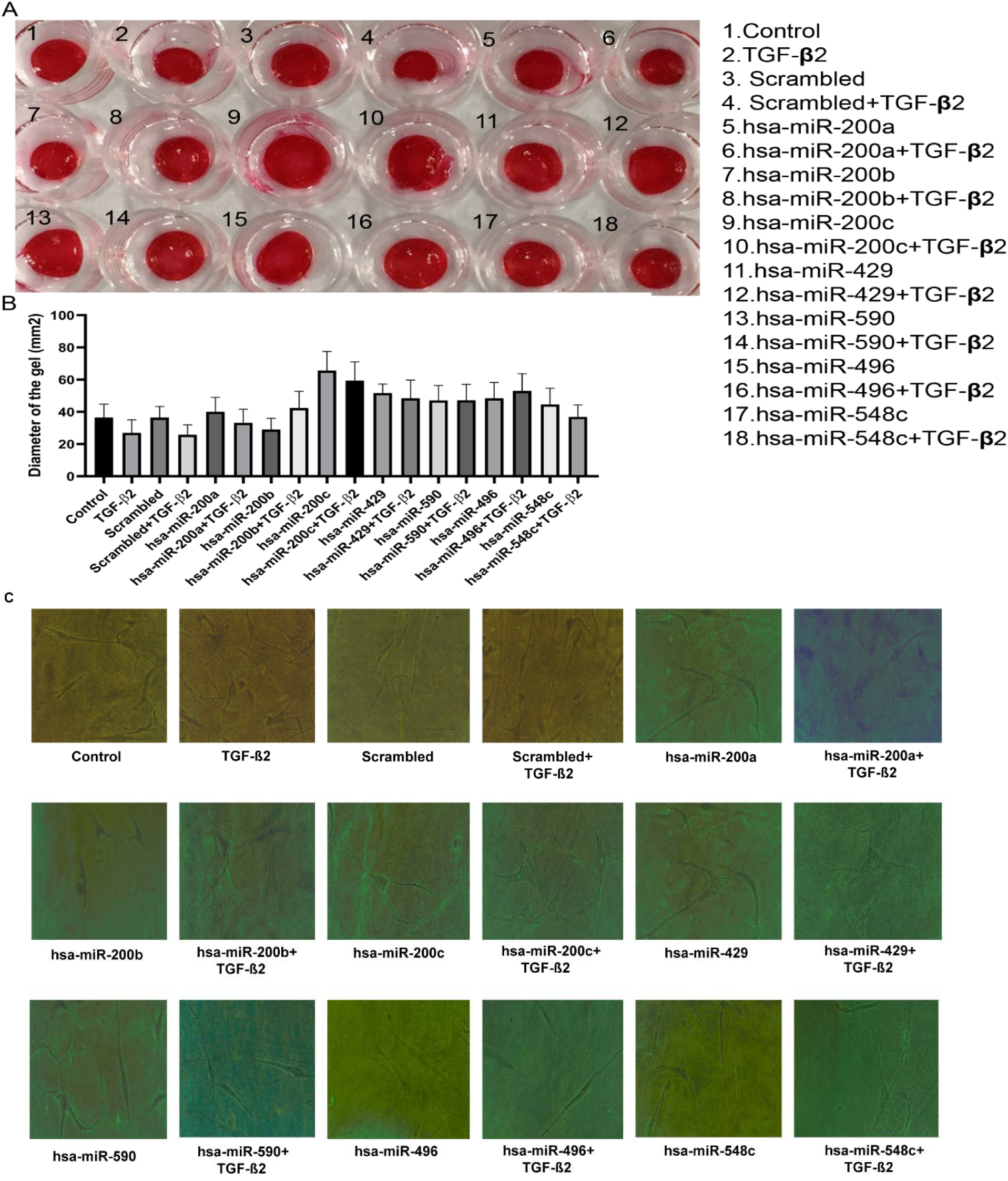
*miR-200* family, *miR-429*, *miR-590*, *miR-496* and *miR-548c* regulated contraction and relaxation of primary HTM cells. (A) Collagen contraction assay with representative collagen gel image of HTM cells transfected with miRNAs or control in combination with TGF-β2 for 48h. (B) Graph representing the area of collagen gel using Image J software. (C) Brightfield images of collagen-embedded cells post-transfection and treatment.

### 2.8 TGF-β2, miRNAs, and IOP follow circadian rhythmic patterns in in-vitro and murine models

Numerous studies have demonstrated that IOP exhibits a circadian rhythmic pattern in human and experimental animal models ^57^. Similarly, the expression of *TGF-*β*2* also displays a circadian pattern, and its over-expression in normal mouse eyes has been shown to alter the rhythmic pattern and elevate IOP ^58^. Our findings suggest that *TGF-* β*2*, miRNA, and glaucoma candidate genes may regulate each other at multiple levels in a feedforward and feedback manner. This observation led us to investigate the expression patterns of selected representative biological clock genes, glaucoma candidate genes, and predicted miRNAs in the trabecular meshwork cells, to elucidate their involvement in IOP regulation.

To achieve this, HTM cells were serum-starved overnight and then replenished with a complete growth medium containing TGF-β2. Cell lysates were collected at 4h intervals, starting from 0h to 48h. Our qRT-PCR analysis revealed that the expression of *BMAL1*, *CLOCK*, *PER1*, *PER2*, *CRY2*, *CRY1*, and *Rev-Erb* α genes follows a circadian rhythmic pattern in primary trabecular meshwork (HTM) cells (Figures 7A-G).

**Figure 7:**
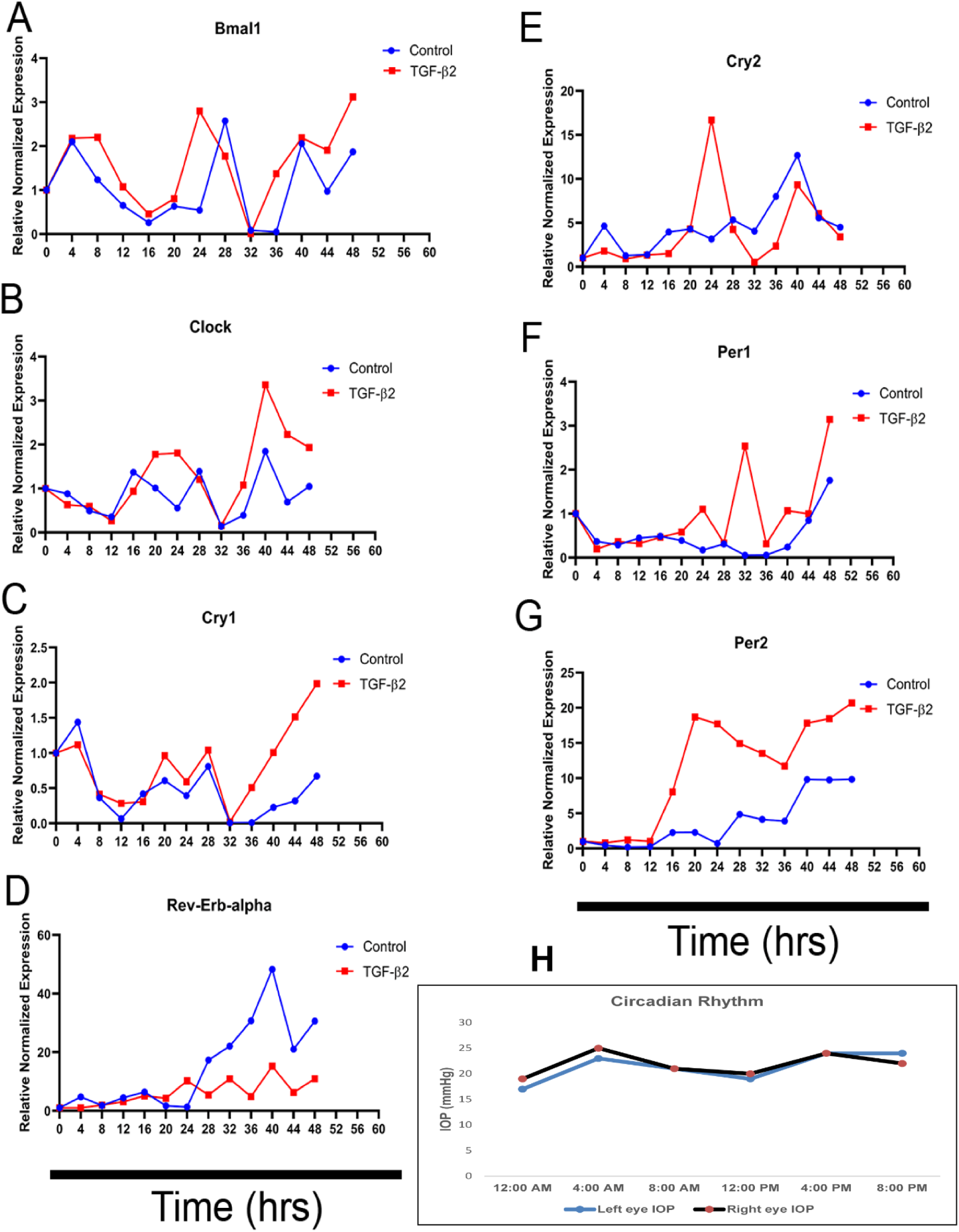
TGF-β2, miRNAs, and IOP follow circadian rhythmic patterns in in-vitro and murine models. (A-G) RT-PCR analysis o with and without TGF-β2 for different time points from 0-48h. (H) IOP (mmHg) measured at different time intervals in 24h cycle post-treatment of dex in C57BL/6J mice.

To check the circadian pattern of IOP and effects of dexamethasone treatments in murine models, one of the eyes of each male C57BL/6 mice was topically treated with 0.1% dexamethasone whereas the other eye was with vehicle control for 3 weeks and measured IOP using rebound tonometer, Icare® TONOLAB at the end of every week until 3 weeks of treatment. Intraocular pressure measurements of control eyes at different time points 12 AM, 4 AM, 8 AM, 12 PM, 4 PM, and 8 PM showed the circadian pattern whereas dexamethasone-treated eyes showed a disturbed pattern with high IOP compared with controls (Figure 8H).

**Figure 8:**
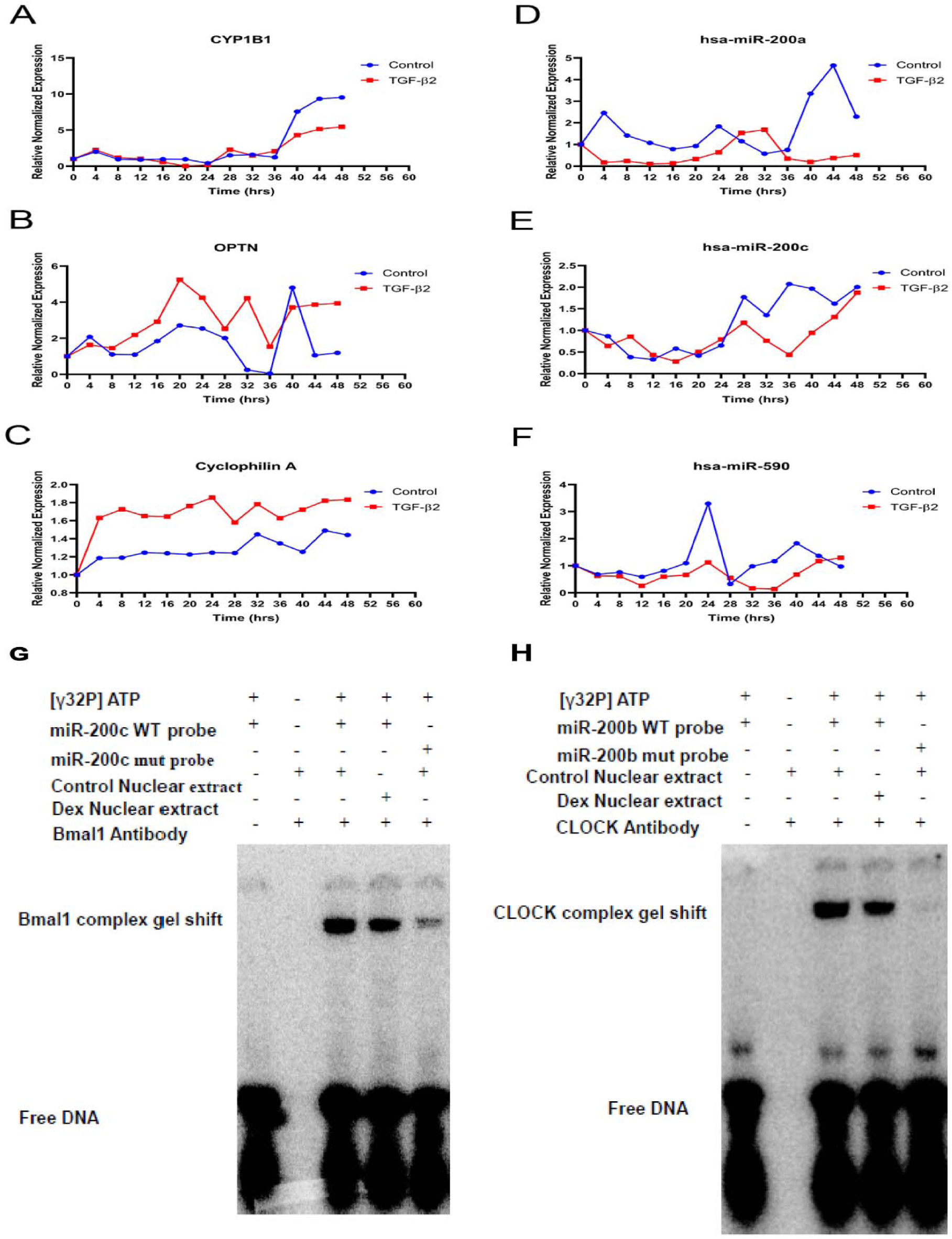
TGF-β2, miRNAs, and IOP follow circadian rhythmic patterns in in-vitro and murine models. (A-F) RT-PCR analysis at different time points between 0-48h. (G, H) EMSA showing binding of BMAL1 on *miR-200c* promoter and CLOCK on *miR-200b* promoter respectively.

As HTM cells exhibit a circadian rhythm, we conducted an analysis of the expression patterns of representative glaucoma candidate genes and miRNAs about circadian rhythmic genes. Serum-starved primary HTM cells were treated with or without TGF-β2 for 0h to 48h and qRT PCR analysis was performed. Our findings revealed that miRNAs (*miR-200a*, *miR-200c*, and *miR-590*), along with the *CYP1B1* and *OPTN* genes, displayed a distinct expression pattern similar to that of clock genes. However, treatment with TGF-β2 treatment (2ng/ml) for 0-48 hours disrupted or delayed the rhythmic expression of *miR-200a*, *miR-200c*, *miR-590*, *CYP1B1*, and *OPTN* at each time point compared to control treatments (Figures 8A-F). Cyclophilin A served as a negative control for gene expression comparison.

To elucidate the mechanistic correlation, we investigated the relationship between clock genes and miRNAs/glaucoma candidate genes. Interestingly, our bioinformatics analysis using the Genomatix tool predicted potential binding sites of CLOCK and BMAL1 to the promoters of *miR-200b* and *miR-200c*, respectively. This binding was further validated by EMSA (Figure 8G-H). In summary, these results suggest that the interplay between TGF-β2 and miRNAs may contribute to the rhythmic regulation of IOP.

## Discussion

Studies investigating the genetic causes of glaucoma etiology have identified loss-of-function mutations in genes such as *CYP1B1, FOXC1, MYOC, OPTN, WDR36,* and *LTBP2*, among others, which are associated with glaucoma. However, the precise molecular mechanisms underlying disease development remain elusive^48,49^. In this study, we uncovered a functional correlation between glaucoma candidate genes. Specifically, we demonstrated that *OPTN* loss downregulates CYP1B1 expression by up-regulating FOXC1 expression in TM cells (Figure 2D). Additionally, we confirmed that FOXC1 negatively regulates *CYP1B1* by binding to the *CYP1B1* promoter sequence (Figure 2F). Moreover, ectopic expression of *OPTN* led to FOXC1 degradation by ubiquitination (Figure 2J). Phenotypically, *OPTN* loss in primary HTM cells increased the formation of actin stress fibers and focal adhesions (Figure 3(II)), thereby resulting in elevated IOP. Our results delineate a functional relationship between OPTN, CYP1B1, and FOXC1 at the molecular level, which may influence cytoskeletal rearrangement in HTM cells, subsequently impacting the regulation of normal IOP.

The cytokine TGF-β2 is highly expressed in the aqueous humor of glaucoma patients and is implicated in various pathological conditions ^20^. Our findings demonstrate that TGF-β2 can upregulate the expression of FOXC1, MYOC, OPTN, and LTBP2 while concurrently downregulating CYP1B1 (Figure 1L-G). Notably, we demonstrate for the first time that the *CYP1B1* promoter contains a consensus FOXC1 binding site (SF2), allowing FOXC1 to negatively regulate *CYP1B1* expression (Figure 2E, F). This observation aligns with our finding that TGF-β2 downregulates CYP1B1 expression by upregulating FOXC1 (Figure 1L). Moreover, while TGF-β2 treatment led to the upregulation of OPTN (Figure 1L, P), overexpression of *OPTN* resulted in the downregulation of TGF-β2. Conversely, *OPTN* knockdown led to the upregulation of TGF-β2 (Figure 3(I)C). These results suggest that *OPTN* may function as a regulatory sensor, exerting its effect through TGF-β2, thereby modulating the expression of *FOXC1* and *CYP1B1*. In summary, these results elucidate the intricate relationship between glaucoma candidate genes and TGF-β2.

We also observed an upregulation of the Rho/ROCK pathway, which is implicated in TM function, in *OPTN* knockdown cells (Figure 3(I)C), thus suggesting a correlation between the activation of Rho/ROCK signaling in primary HTM cells and its activation in glaucomatous patients. This correlation underscores a potential interplay between glaucoma candidate genes and TGF-β2, which subsequently influences the formation of actin stress fibers and focal adhesion (Figure 3(I)). These cellular changes might contribute to outflow resistance and elevated IOP in the anterior chamber of the eye.

Long-term glucocorticoid treatment has been linked to elevated IOP and glaucoma development ^44^. Our findings demonstrate that treatment with the glucocorticoids dexamethasone and prednisolone induces the upregulation of FOXC1, MYOC, and OPTN while downregulating CYP1B1 (Figure 1A-F). Additionally, we observed activation of the Rho/ROCK pathway and increased formation of actin stress fibers and focal adhesions in primary TM cells, mirroring the effects of TGF-β2 treatment (Figures 3(I)). These observations suggest a potential correlation between the pathological conditions observed in glaucomatous TM tissue ^6,20^. Furthermore, glucocorticoid treatment increased TGF-β2 levels in TM cells (Figure 1G, H), suggesting a potential mechanism through the TGF-β2 signaling pathway. Consequently, our results imply that TGF-β2 signaling might mediate the effects of dexamethasone. Indeed, our observations indicate that the activation of TGF-β2 and Rho/ROCK signaling pathways leads to aberrant signaling patterns that mimic glaucoma development. This activation promotes the contraction of TM tissue, resulting in increased outflow resistance and elevated IOP in the anterior chamber of the eye.

Clinical studies have highlighted the efficacy of ROCK inhibitors, such as (Y-27632), in reducing IOP in glaucoma patients ^52^, implying a potential involvement of intrinsic ROCK regulators, including miRNAs and epigenetic factors, in mediating these effects within TM cells. Using bioinformatics tools, we predicted that certain miRNAs could modulate TGF-β2 and ROCK2 expression in primary HTM cells. Notably existing literature demonstrated that *miR-200a*, *miR-200b*, *miR-200c*, *miR-429*, and *miR-141* exhibit unique expression patterns in trabecular meshwork cells compared to ciliary epithelium, underscoring their importance in the outflow pathway of the aqueous humor. This suggests a potential link between these miRNAs, TGF-β2, ROCK2, and the regulation of IOP, thereby enhancing our understanding of the molecular mechanisms underlying glaucoma pathogenesis.

Furthermore, we discovered that the ectopic expression of these miRNAs led to the downregulation of Rho/ROCK signaling, suppression of actin filament polymerization, reduction of focal adhesions, and relaxation of collagen in primary HTM cells, even after TGF-β2 treatment. Also, TGF-β2 downregulated these miRNAs (SF5A) in primary HTM cells, illustrating the interplay between TGF-β2 and these miRNAs. Additionally, ectopic expression of putative miRNAs resulted in the downregulation of *FOXC1 and CYP1B1* expression. Conversely, *miR-200a*, *miR-200b*, *miR-200c*, *miR-429*, *miR-141,* and *miR-548c* upregulated *OPTN* expression in primary HTM cells, whereas this effect was not observed for *miR-590* (Figure 4B). These findings highlight the intricate interplay between candidate genes (*FOXC1, CYP1B1, and OPTN)* with TGF-β2 and the predicted miRNAs in TM cells, providing further insights into the molecular mechanisms underlying glaucoma pathogenesis.

In the normal physiological life cycle, the production and flow of aqueous humor follow a 24-hour cycle, peaking during the morning (diurnal) and declining at night (nocturnal). This cycle correlates with lower IOP in the morning and higher IOP at night ^11^. Our analysis revealed that the expression patterns of clock genes, including *BMAL1, CLOCK, PER1, PER2, CRY2, CRY1, and Rev-Erb* α as well as *TGF-*β*2*, *CYP1B1, OPTN, miR-200a*, *miR-200c*, *miR-590* were synchronized in primary HTM cells, consistent with circadian rhythmic patterns (Figure 7,8). Interestingly, previous studies have demonstrated the ability of FOXC1 to bind to the *CLOCK* promoter, modulating its expression ^40^. Through in silico analysis and EMSA studies, we identified a consensus binding site for the BMAL1 protein in the *miR-200c* promoter, as well as a consensus sequence for the CLOCK protein in the *miR-200b* promoter (SF6), thus supporting the interplay between circadian CLOCK mechanisms and gene expression regulation in TM cells. Hence, our findings emphasize the potential regulatory role of circadian rhythms in the molecular pathways involved in IOP and glaucoma development.

Research has shown that OPTN inhibits TGF-β signaling in TNBC (Triple negative breast cancer) cells by interacting with the TGF-β type I receptor (TβRI), leading to its ubiquitination and degradation. This mechanism suppresses metastasis by reducing EMT-associated factors ^59^ experimental findings in TM cells indicate that *OPTN* overexpression suppresses TGF-β signaling, whereas *OPTN* knockdown enhances TGF-β signaling and EMT. Furthermore, *OPTN* overexpression leads to FOXC1 ubiquitination, a process reversed by proteasomal inhibitors. These findings suggest that OPTN may modulate the balance between diurnal vs nocturnal IOP by regulating various key signal intermediaries, thereby contributing to the regulation of IOP fluctuations over the 24-hour cycle.

In conclusion, our study highlights the intricate regulatory network involving TGF-β2, the *miRNA-200* family, and key genes such as *OPTN*, *FOXC1,* and *CYP1B1*, which operate in concordance with circadian rhythm genes. Perturbations among these regulatory factors are associated with fluctuations in nocturnal and diurnal IOP. Disruption of this regulatory pathway due to mutations, altered metabolism, or disturbances in the CLOCK cycle can lead to aberrant aqueous humor dynamics and IOP, ultimately contributing to glaucoma pathogenesis.

## Limitations of the study

1. The study has not discussed the molecular mechanism of how TGF-β2 inhibits the levels of putative miRNAs.
2. Study lacks clinical data however the data is correlating the clinical observations.
3. The study is limited to the mechanism operating in the anterior segment of the eye.

## STAR Methods

### *In silico* miRNA target prediction

miRNA target prediction was performed using open-sourced bioinformatics algorithm tools based on sequence complementarity, such as TargetScan8.0 (http://www.targetscan.org), miRanda (http://www.microrna.org) and miRBase (http://www.mirbase.org/), for all putative candidate genes in the present study (*CYP1B1*, *MYOC*, *OPTN*, *FOXC1*, *LTBP2*, *TGF-*β*2* and *ROCK2*). Predicted miRNAs were listed in the (Table 1)

### Cell culture

Primary human trabecular meshwork (HTM) cells and its complete culture medium with growth factors (TMCM kit) were purchased from ScienCell Research Laboratories Inc. and cultured in the trabecular meshwork cell medium (TMCM) medium, (2%) fetal bovine serum (FBS) and (1%) trabecular meshwork cell growth supplement (TMCGS), (1%) penicillin-streptomycin (ScienCell Research Laboratories Inc., Carlsbad, CA), incubated at 37°C in 5% CO_2_ incubator with 95% air. Recombinant human TGF-β2 (2ng/ml; Sigma-Aldrich), dexamethasone (300nM/ml; Sigma) and prednisolone (300nM/ml; Sigma) were used. miRNA mimics and anti-miR mimics against each respective miRNA were purchased from (Qiagen) (Table 2) and were transfected in primary HTM cells using Dharmafect duo transfection reagent (Dharmacon). HEK 293T cell lines (ATCC) cultured in DMEM (Invitrogen-Gibco Life Technologies) were used for luciferase studies.

### RNA isolation and quantitative real-time PCR (qRT-PCR)

Total RNA was extracted with RNAiso Plus (Takara) for genes and miRNeasy isolation kit (Qiagen) for miRNAs following the manufacturer’s protocols. Real-time PCR was performed by iTaq^TM^ Universal SYBR^®^ Green Supermix (Bio-Rad) for the expression of genes and QuantiTect SyBr green PCR system (Qiagen) for the expression of miRNAs on a CFX96 (Bio-Rad) in triplicates on Complementary DNA (cDNA) synthesized from total RNA using iScript^TM^ cDNA synthesis kit (Bio-Rad) for genes and miScript II RT Kit (Qiagen) for miRNAs. Data were collected and analyzed by ddCT method. Data values were normalized to GAPDH for data analysis and quantification of genes. For miRNA analysis, real-time PCR was performed using miRNA prime assays (Qiagen), and U6 small nuclear RNA (snRNA) was used for normalization.

### Transfection of Si-RNAs, synthetic microRNA, and its inhibitors

Primary human trabecular meshwork (HTM) cells were seeded (1.5 × 10^5^ cells) per well in 6 well plates and transfected with synthetic miRNA mimics (Qiagen) at a final concentration (80 nM) using DharmaFECT^TM^ duo transfection reagent (Dharmacon, GE Healthcare) in Opti-MEM^TM^ (Invitrogen-GIBCO Life Technologies) and replenished with TMCM complete medium after transfection. Total RNA and protein were collected at 48 hours post-transfection for analysis. Whereas miRNA inhibitors, (Anti-miRs, Qiagen) were transfected at a final concentration (250 nM) against each respective miRNA or a negative control inhibitor. For experiments where CYP1B1, FOXC1, and OPTN were concurrently knocked down, 100pM of each siRNA (Qiagen) or a control siRNA (All-star negative control, Qiagen) were used.

### Western blotting

Transfected and treated HTM cells, and made cell lysates in (1x) radioimmunoprecipitation assay (RIPA) lysis buffer (Cell Signaling Technology, CST) containing protease inhibitor cocktail (Pierce, Rockford, IL) were fractionated on (10%) SDS polyacrylamide gel followed by transfer onto a nitrocellulose membrane, (Bio-Rad laboratories). Blots were probed with anti-CYP1B1, anti-OPTN, anti-MYOC, anti-LTBP2, anti-p-Cofilin, anti–Cofilin, anti-Paxillin, anti-ROCK2, anti-TGF-β2 and anti-GAPDH (Santa Cruz Biotechnology) and anti-FOXC1, anti-CLOCK, anti-BMAL1 (CST) antibodies (Table 3) followed by respective secondary antibodies (Table 4) (CST) and immunocomplexes were detected using the electrochemiluminescence, ECL method (GE Healthcare).

### Immunofluorescence Staining

Primary HTM cells plated onto poly-L-lysine (Sigma) coated coverslips were transfected with miRNA mimics followed by treatment with recombinant human TGF-β2 (2ng/ml; Sigma) as described above and subjected to staining, 24 hours post-transfection. Cells were fixed in (4%) paraformaldehyde, permeabilized with (0.1%) Triton X-100, and probed with a rabbit-anti-paxillin antibody (1:500; Santa Cruz Biotechnology) followed by goat-anti-rabbit-Alexa flour 488 conjugated antibodies (1:500; Invitrogen) (Table 4).

For F-actin staining, fixed and permeabilized cells were incubated with rhodamine-phalloidin (Invitrogen) for 10 min followed by mounting along with 4’-6-Diamidino-2-phenylindole (DAPI; Invitrogen). Images were captured using a Leica TCS SP8 confocal microscope. Images were analyzed with Leica Cell software.

### 3’UTR reporter analysis

The 3’UTRs of *TGF-*β*2*, *ROCK2*, *CYP1B1*, *FOXC1*, and *OPTN* were amplified by PCR from genomic DNA isolated from HEK293T cell lines, cloned into the pMIR-REPORT luciferase expression vector (Thermo Fisher Scientific). Luciferase reporter plasmid (100ng) and pRL-TK control (200ng for normalization; Addgene) were transfected with dharmafect duo (Dharmacon, GE Healthcare) transfection reagent into HEK293T cell lines seeded in 24 well plates (6×10^4^ cells per well). For co-transfection experiments, 5 nM of synthetic miRNAs (pre-miRs, Qiagen) or (30 nM) of miRNA inhibitors (Anti-miRs, Qiagen) were added to the above reactions. Cells were collected after 48h of post-transfection and subjected to dual-luciferase reporter assay (Promega) as described in product protocol. All experiments were performed in triplicates for at least three independent experiments.

### Enzyme-Linked Immunosorbent Assay (ELISA)

Conditional Medium was collected from the samples and TGF-β2 levels were quantified using an hTGFβ2 ELISA kit purchased from R&D Systems, according to the manufacturer protocol. Briefly, Supernatants collected from the cells were subjected to centrifugation to remove cell debris. Samples were activated by subjecting them to acidification. Further steps were followed as per the protocol and the optical density of the samples at 450 nm was measured using plate reader molecular devices spectra max M2^e^. For wavelength correction, all the samples were measured at 570 nm. Calculations were done by plotting a four-parameter logistics (4-PL) curve fit. The concentration read from the standard curve multiplied by the dilution factor, 7.8.

### *In-vivo* dexamethasone treatments and IOP measurements

C57BL/6J mice (∼3 months old male) were purchased from Liveon Biolabs Private Limited after obtaining Institutional animal-ethical clearance. All the procedures and protocols were followed as per the approvals by the ARRIVE (Animal Research Reporting- In Vivo Experiments-mentioned in journal guidelines and vision research). Topical administration of dexamethasone phosphate (0.1%, Sigma) was given to the right eye of the mice and sterile phosphate-buffered saline (PBS) was given to the left eye as a control daily 3 times for up to 3 weeks. Doses were given daily between 9-10 am; 1 to 2 pm; and 6 to 7 pm. Mice were loosely held for a few seconds after the application of eye drops to ensure effective penetration into the eye. At 3 weeks IOP was measured using a rebound tonometer (TONOLAB, ICARE Instruments), as per the instructions of the manufacturer at every 4h interval period for a complete 24h cycle precisely at 0, 4, 8, 12, 16, and 20 h. At the end of the study, whole eyeballs were fixed in 10% formaldehyde (Sigma) and sectioned for immunohistochemistry.

### Electrophoretic mobility shift assays (EMSA)

Putative transcription factor binding sites were predicted using Genomatix: Matinspetor tool (https://www.genomatix.de/online_help/help_matinspector/matinspector_help.html) for identification of transcription factor binding and consensus sequencing on promoter regions. Confluent primary HTM cells in a 100mm dish were treated and nuclear extracts were collected with nuclear extraction buffer containing 20 mM HEPES (pH-7.4), 0.4mM NaCl, 1mM EDTA (pH-8), 0.1M EGTA and protease inhibitor cocktail. The 5’ end of CYP1B1, *miR-200b,* and *miR-200c* promoter sequence along with mutant probes was end-labeled with 20 fmol of [γ32P], and the unlabeled probe was removed with illustra Microspin-G-50 columns. Nuclear fractions were incubated with and without FOXC1/CLOCK/BMAL1 antibody for the promoters of CYP1B1, *miR-200b* and *miR-200c* probe respectively, and 10pmol of probe at 37°C for 30 min in 1X binding buffer (200mM HEPES (pH 7.4), 40mM DTT, 4mM EDTA (pH-8.0) and 50% glycerol). The nuclear protein complexes were loaded on 6% non-denature PAGE gel with 6X DNA loading dye. DNA mobility and band shifts were observed in the phosphor imager system (Bio-Rad). FOXC1, CLOCK, and BMAL1 antibodies were procured from cell signaling technologies (Table 3).

### Oligonucleotide sequences

All real-time PCR primers of CYP1B1, FOXC1, OPTN, MYOC, LTBP2, TGF-β2, CLOCK, BMAL1, PER1, PER2, CRY1, CRY2, Rev-Erbα, GAPDH and 3’UTRs primers of TGF-β2, ROCK2, CYP1B1, OPTN and FOXC1 and EMSA primers of CYP1B1, *miR-200b* and *miR-200c* promoter sequences are given in (Table 4).

### Statistical Analysis

The data values were expressed as mean ± SEM statistical analysis was performed using the student t-test. The differences between the mean values of control and treated/ transfected groups were analyzed using student t-test. P-value lesser than 0.05 was considered statistically significant. For all cell culture studies, a minimum of three independent experiments were carried out.

## Author contributions

AB: Conceived, conceptualized the idea, and compiled the information. CKS & NNG conceptualized idea, executed experiments, compiled and written manuscript SC: executed experiments, critical comments. All authors reviewed the manuscript.

## Acknowledgements

The authors acknowledge the University of Hyderabad for funding the laboratory through institutional funding from DST-FIST-Level II, the Institute of Eminence (IoE) (UoH-IoE-IIRC-23-001), and DBT-SAHAJ/BUILDER (BT/INF/22/SP41176/2020). ABMR acknowledges support from the DST-SERB (EMR/2016/007595), DBT-RNAi/GET (BT/PR8805/AGR/36/747 and BT/PR38397/GET/119/306/2020) governments of India through an independent extramural grant. CKS and SC acknowledge UGC-CSIR and NNG to ICMR-SRF for fellowship for doctoral studies.

