## Supplemental Figures for "TGF-β2/OPTN/FOXC1/*miR-200* axis regulates intraocular pressure dynamics in trabecular meshwork cells"

**Supplementary Figures (SF)**

**A**

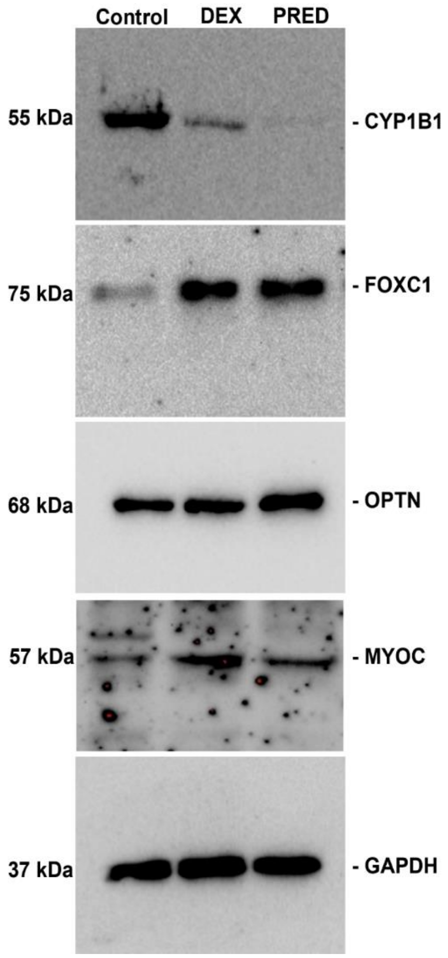


## SF1:

(A) Original western blot images corresponding to Figure 1A.

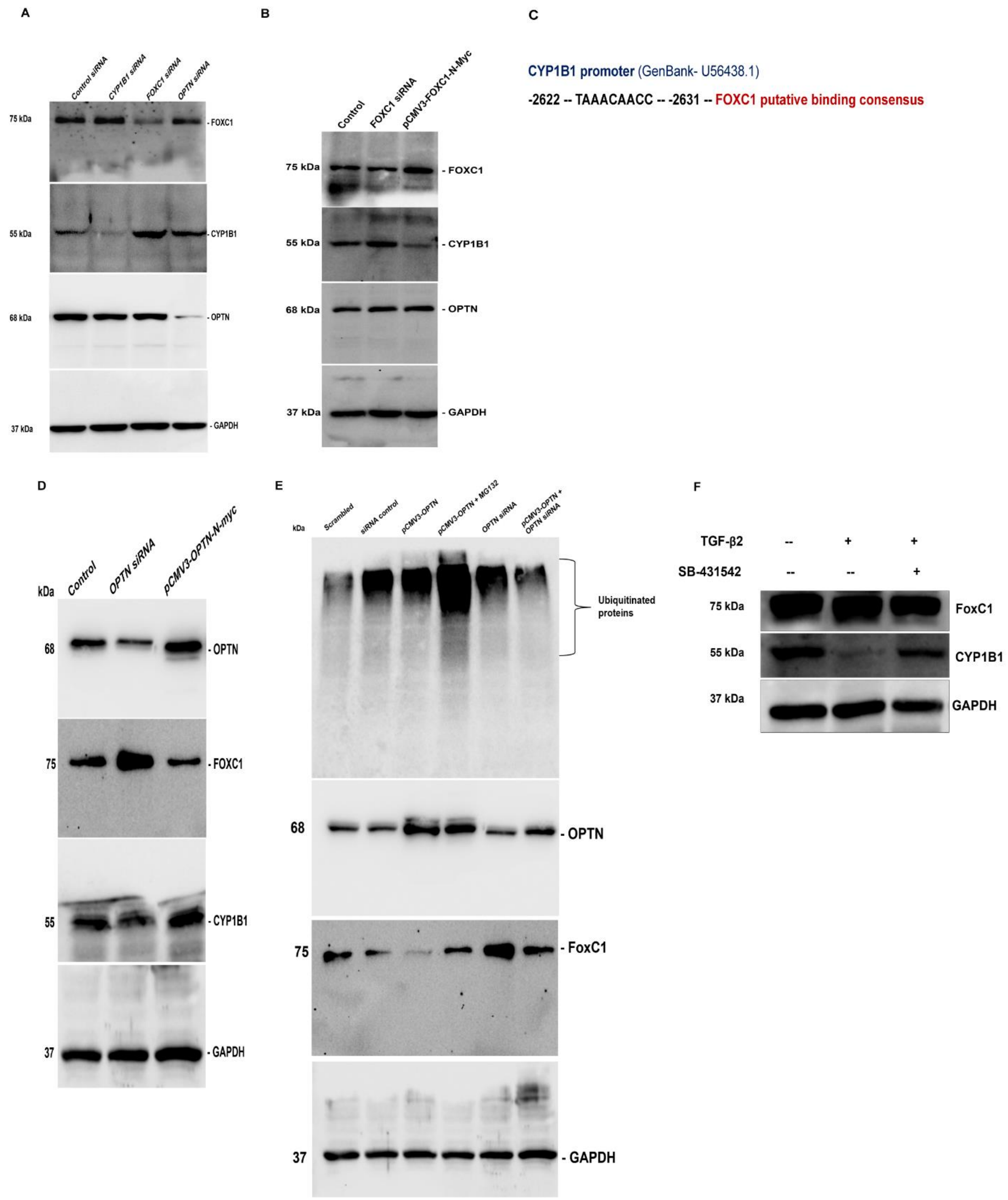


**SF2:**

(A) Original western blot images corresponding to Figure 2D.
(B) Original western blot images corresponding to Figure 2E.

(C) Schematic representation of FOXC1 putative binding consensus on *CYP1B1* promoter.

(D) Original western blot images corresponding to Figure 2H.

(E) Original western blot images corresponding to Figure 2J.

(F) Western blot analysis in HTM cells 48h post treatment (24h treatment with SB-431542, TGF-β pathway inhibitor followed by TGF-β2 treatment for 24h).

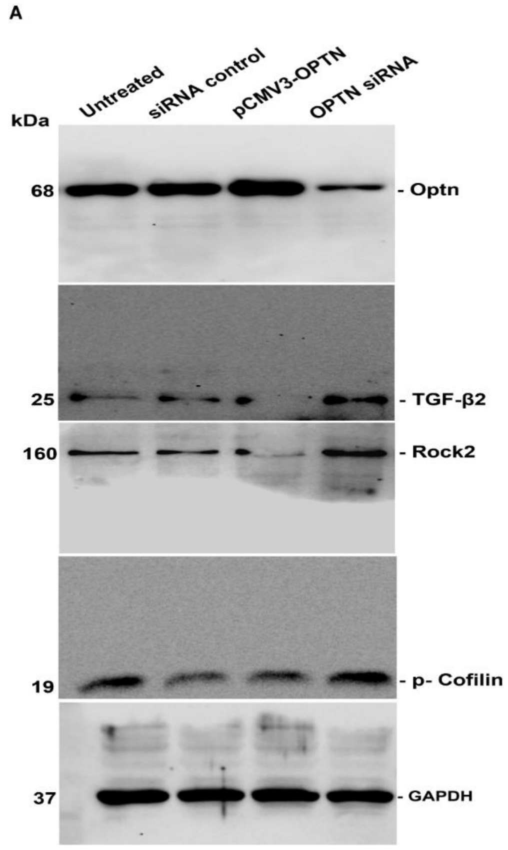


## SF3:

(A) Original western blot images corresponding to Figure 3(I)C.

A

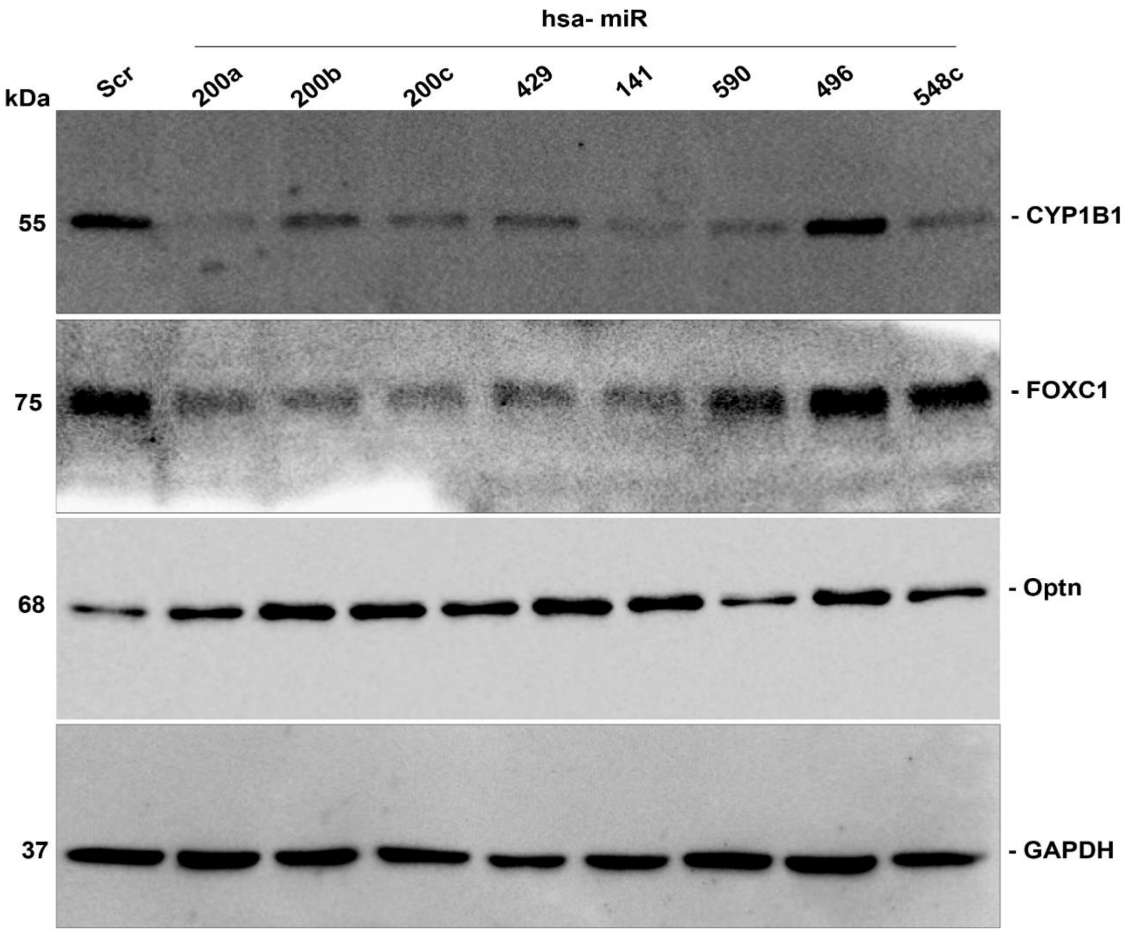

hsa- miR
kDa
Scr
200a
200b
200c
429
141
590
496
548c
55
75
68
37
- CYP1B1
- FOXC1
- Optn
- GAPDH


B

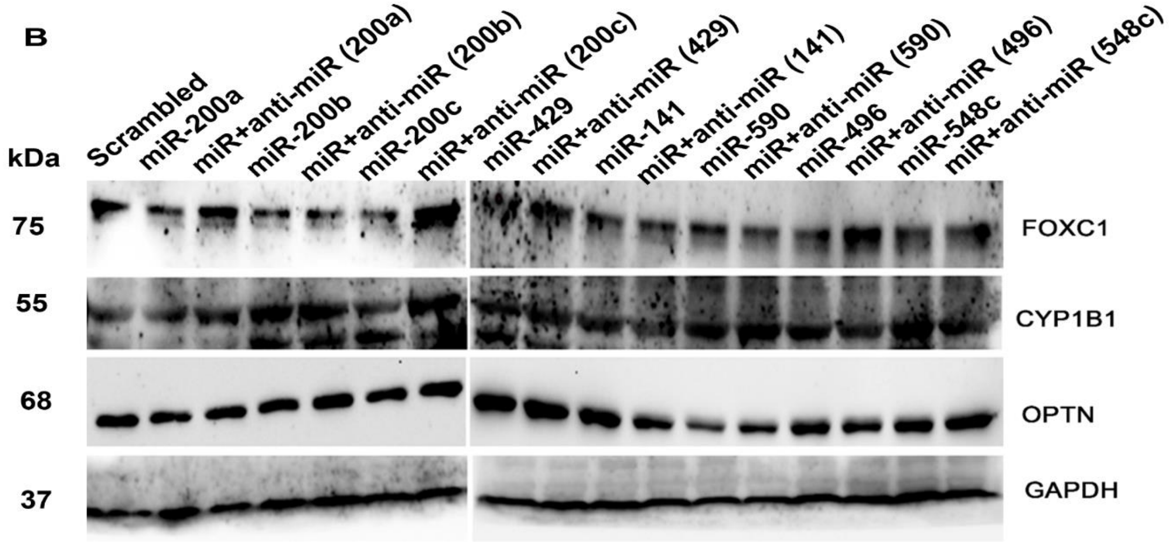

kDa
Scrambled
miR-200a
miR+anti-miR (200a)
miR-200b
miR+anti-miR (200b)
miR-200c
miR+anti-miR (200c)
miR-429
miR+anti-miR (429)
miR-141
miR+anti-miR (141)
miR-590
miR+anti-miR (590)
miR-496
miR+anti-miR (496)
miR-548c
miR+anti-miR (548c)
75
55
68
37
FOXC1
CYP1B1
OPTN
GAPDH


SF4

**SF4:**

(A) Original western blot images corresponding to Figure 4B.

(B) Western blot image corresponding to Figure 4(C-E).

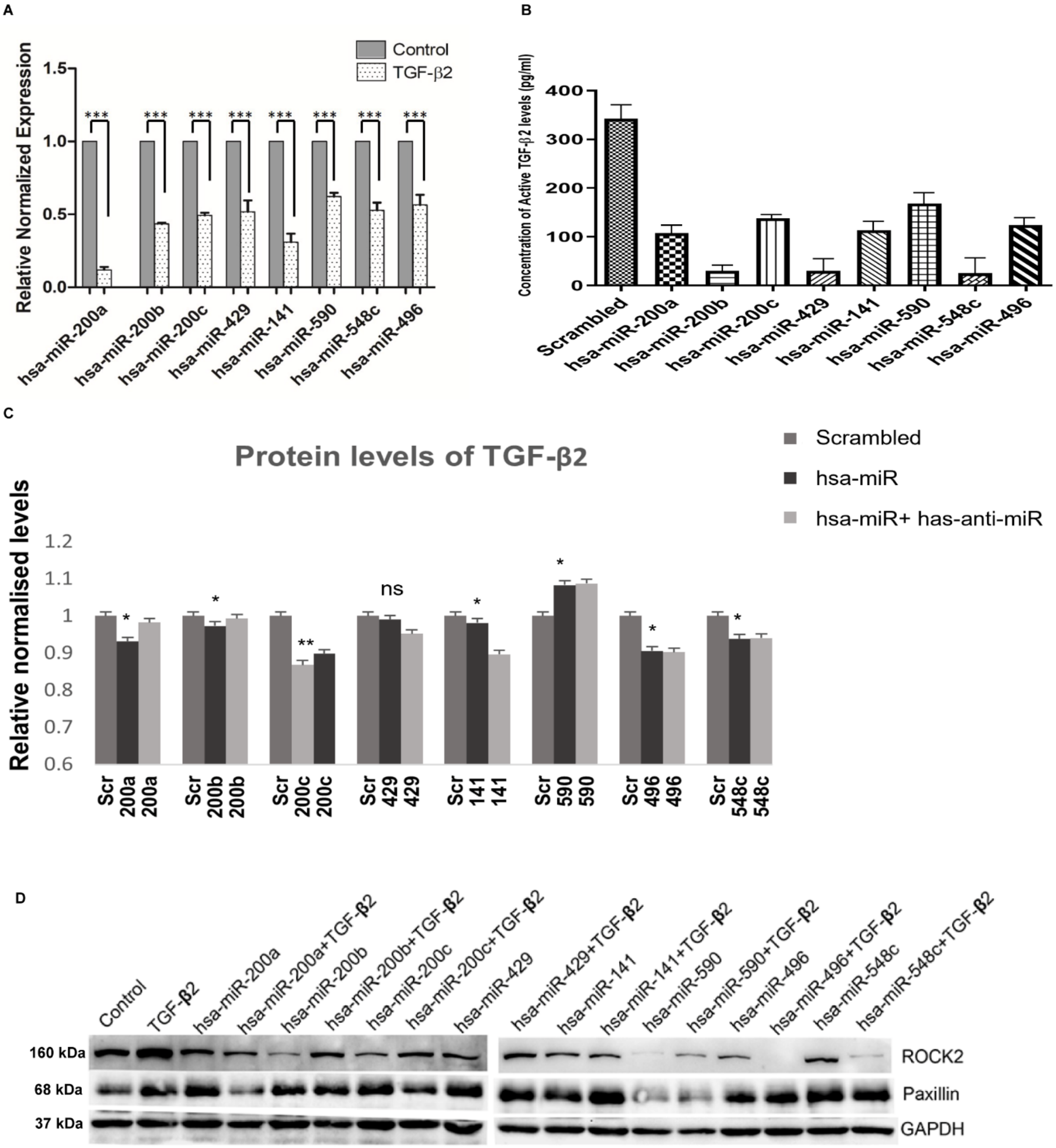


**SF5**

**SF5:**

(A) RT-PCR analysis of miRNAs (miR-200a, miR-200b, miR-200c, miR-429, miR-141, miR-590, miR-548c, miR-496) expression upon treating HTM cells with TGF- β2 for 48h.

(B) ELISA showing active concentration of TGF-β2 in the conditioned medium 48h post transfection of miRNAs (miR-200a, miR-200b, miR-200c, miR-429, miR-141, miR-590, miR-548c, miR-496).

(C) Western blot quantification data of protein lysates upon transfection of respective miRs and anti-miRs corresponding to TGF-β2.

(D) Western blot analysis upon transfecting HTM cells with respective miRNAs in combination with treatment of TGF-β2.

**A**

**miR-200c promoter**

**-932 -- aggagttaTGTGcagcc -- -948 -- BMAL1 putative binding consensus**

**-940 -- anchoring position**

**B**

**miR-200b promoter**

**+2501-- aggggtCACGttcttgc -- +2517 -- CLOCK putative binding consensus**

**+2509 -- anchoring position**

Note: +/- in the position implies strand

**SF6:**

(A) Schematic representation of BMAL1 putative binding consensus on *miR-200c* promoter.

(B) Schematic representation of CLOCK putative binding consensus on *miR-200b* promoter.
